# What drives mixed-species shoaling among wild zebrafish? Role of predators, food access, abundance of conspecifics and kin familiarity

**DOI:** 10.1101/2022.07.11.499549

**Authors:** Ishani Mukherjee, Anuradha Bhat

## Abstract

Mixed-species groups commonly occur across a wide range of faunal communities and are known to provide several benefits to members. While zebrafish have often been observed to form mixed-species shoals with coexisting species, the factors determining their occurrence is not yet understood. Using laboratory-based experiments, we decipher the main ecological drivers of mixed-species shoaling in these tropical fish communities. Shoals comprising zebrafish (*Danio rerio*), flying barbs (*Esomus danricus*) and whitespots (*Aplocheilus panchax*) were collected from a stagnant canal at Haringhata (West Bengal, India). Experiments to assess foraging efficiency were conducted where single or mixed-species shoals (comprising 5 individuals) were given low or high amounts of food. Shoal choice experiments were also conducted to assess the preferences of test subjects (zebrafish individuals) for forming associations based on shoal composition and familiarity. Results from experiments on feeding efficiency revealed that foraging time varied substantially among the shoal types (i.e., single or mixed-species), and was dependent on the quantity of food available, but not linked to the body size of species composing the shoal. The choice experiments to examine preference for associations revealed that under predator risk, zebrafish associate more with mixed shoals, and showed comparable associations to shoals differing in the abundance of conspecifics. Furthermore, we found that zebrafish preferred to associate with familiar conspecific over unfamiliar mixed and unfamiliar conspecific shoals. Therefore, equitable food consumption in mixed shoals, greater association to mixed shoals in presence of predator and familiarity were found to be important drivers for choosing mixed-species shoaling by zebrafish.

## Introduction

Predator avoidance and the need to forage efficiently are the main drivers of group living across a wide range of taxa (Rubenstein 1978). The cohesion and longevity of group formation in organisms vary based on a variety of factors like sex-composition, predation pressure and nutrient availability (Bon et al. 1990; Ward and Webster 2016; Herbert-Read et al. 2017; Iaonnou 2021). These can also further depend on age, species composition, and personality types of members within these groups (Ward et al. 2002; Goodale et al. 2020). Mixed-species groups comprise individuals belonging to more than one species, in spatial proximity to each other and interacting with one another to maintain the group (Goodale et al. 2017). Such groups often confer more benefits as compared to single species groups of comparable sizes. (Goodale et al. 2017; Stensland et al. 2003). Multi-species biofilms comprising many different species of microorganisms have greater resistance to certain antimicrobials as compared to the ones comprising a single species (Lee et al. 2014). Several bird species form multi-species flocks as these provide group members with enhanced anti-predator and foraging benefits (Hino 2003; Sridhar et al. 2009; Sridhar et al. 2013). Mixed-species associations have been observed involving two monkey species *(Cercopithecus diana* and *Cercopithecus campbelli)*, where both species benefit from the association in terms of improved foraging and increased social behavior (Wolters and Zuberbühler 2003). Zebrafish are also known to form mixed shoals with a variety co-occurring species (Miller and Gerlai 2011; Spence et al. 2008; Suriyampola et al. 2016). In this study, using laboratory-based experiments, we aim to explore and decipher the likely important drivers of mixed-species shoaling comprising wild zebrafish.

Most studies related to shoaling in fishes focus on single species shoals and few consider mixed shoals (Paijmans et al. 2019). Despite this, there are some interesting findings so far: while certain species increase food intake by forming mixed-species shoals, others form mixed shoals to avoid a predator (Lukoschek and McCormick 2000; Mathis and Chivers 2003). The Indo-pacific Sergeant fish (*Abudefduf vaigiensi*) form mixed-species groups to obtain both these benefits i.e., to reduce the chances of predation as well as to increase food access (Paijmans et al. 2020). Mixed-species shoals comprising two species of sticklebacks (three spined stickleback: *Gasterosteus aculeatus* and nine spined stickleback: *Pungitius pungitius*) are as cohesive as shoals of either stickleback species (Ward et al. 2018). Mixed-species shoals are also formed based on familiarity, lack of conspecifics, advantages related to social learning and passive mechanisms such as comparable swimming speed across species (Krause et al. 2005; Paijmans et al. 2019).

Zebrafish (*Danio rerio*) has been a popular model organism for studies on developmental biology, genetics and cell biology. More recently, the behavior and ecology of wild zebrafish has also been widely studied (Engeszer et al. 2007; Spence et al. 2008). Wild zebrafish are found in freshwater streams and ditches across India, Bangladesh and Nepal and a variety of shoal sizes (2-300 individuals) have been reported in the wild (Pritchard et al. 2001; Suriyampola et al. 2016). Wild zebrafish have been reported to shoal with many co-occurring species in these habitats such as flying barbs (*Esomus danricus*), whitespots (*Aplocheilus panchax*), *Rasbora daniconius* and *Puntius* spp (Miller and Gerlai 2011; Spence et al. 2008; Suriyampola et al. 2016). Even as mixed-species shoals are common across a range of zebrafish habitats, shoal properties can vary between populations based on ecological factors like water flow, vegetation and predation pressure (Suriyampola et al. 2016). In this study, we attempt to unravel the primary causes for formation of such mixed-species associations comprising wild zebrafish.

In their natural habitats (comprising of shallow stagnant waterbodies with moderate vegetation) located at Haringhata, West Bengal, India, we observed mixed shoals comprising *Danio rerio, Esomus danricus* and *Aplocheilus panchax*. The mixed shoals were collected, brought to the laboratory and we performed a range of experiments to reveal the drivers of mixed shoaling among wild zebrafish. More specifically, we investigated whether zebrafish is driven towards mixed shoaling due to benefits related to (1) foraging, (2) predator avoidance, (3) lack of conspecifics and (4) familiarity. The rationale for considering improved foraging as a potential driver towards mixed-species shoaling is as follows: It has been established that nutritional state of conspecifics drive shoaling decisions and foraging success of zebrafish shoals (Krause et al., 1999) Within mixed shoals, food might be easily accessible to zebrafish as the other species are known to not show aggression towards heterospecifics (Daniels 2002). Thus, foraging benefits within mixed shoals might be a potential driver towards mixed shoaling in zebrafish. Another factor that controls shoaling in zebrafish is predation (Miller and Gerlai 2011; Speedie and Gerlai 2007) and their natural habitat, zebrafish are prone to predation by the snakehead (*Channa* spp.). Shoaling in mixed-species shoals can be a strategy to evade predation as fish predator’s often selectively prey on a particular species within such mixed shoals (Mathis and Chivers 2003). Here, we hypothesize that *Channa* individuals might have a preference for the larger fishes (whitespots or flying barbs) within the same shoal and this might drive zebrafish towards mixed-species shoaling. Additional factors such as the low abundance of conspecifics, and possible familiarity of heterospecific individuals that share habitats could also drive mixed-species shoaling. We predict that to adhere to ‘safety in numbers’ strategy, zebrafish might shoal with other species when there is low abundance of conspecifics in the habitat. Familiarity is an important driver of inter-shoal associations and field-based studies have revealed that individuals form persistent associations with familiar individuals (Hay and McKinnel 2002; Croft et al. 2004). Further, studies have shown that fish prefer shoaling with familiar heterospecifics over unfamiliar conspecifics (Ward et al. 2003) Therefore, we predict that zebrafish may form mixed shoals based on familiarity with other co-occurring species. We hypothesized that foraging and anti-predator benefits, familiarity with co-occurring species and low relative abundance of conspecifics are the major drivers for zebrafish choosing to form mixed-species shoals. We conducted foraging efficiency and choice-based experiments on wild-caught mixed-species groups of fishes to disentangle the roles of these likely key drivers of mixed-species shoaling.

## Methods

### Collection and maintenance of shoals

This study was conducted on a wild caught zebrafish population collected from its natural habitat at Haringhata (Nadia District) in West Bengal, India. Mixed shoals comprising mostly zebrafish (*Danio rerio)* and flying barbs (*Esomus danricus*) have been observed to be feeding and swimming together at the surface of a stagnant ephemeral ditch. Some shoals also comprise a few whitespots (*Aplochelius panchax*) or juveniles of *Puntius sp*. Though owing to high turbidity and moderate vegetation cover it was not feasible to quantify the exact shoal size and composition, qualitatively, these shoals typically comprised 5-20 individuals, mostly zebrafish and flying barbs, and sometimes whitespots as well. Shoals consisting of zebrafish, flying barbs, and some whitespots were collected using a drag net. The fishes were brought to the laboratory in aerated plastic bags and housed in mixed-species groups of 50 individuals per 60cm × 30cm × 30cm glass aquarium tanks, supplied with aerators. The relative abundance of species in each tank was comparable to the ratio in which they occurred in the natural habitat i.e., each glass tank housed 20 zebrafish, 20 flying barbs and 10 whitespots. Single species tanks (consisting of ∼ 50 individuals of the same species housed together) were also maintained separately and these individuals were used in experiments which involved single species. A temperature range of 24-26°C and a constant lighting condition of 12h dark:12h light was maintained in the holding room. Fishes were fed daily with freeze-dried bloodworms or brine shrimp *(Artemia sp*.). Experiments commenced after a 30-day acclimatization to laboratory conditions.

Snakeheads (*Channa* spp.) are common predators to these fishes and occur widely in the same habitats (Spence et al. 2008). *Channa punctatus* individuals (approximately 16-17.5cm in length) were collected from the same ditch and housed separately in 60 × 30 × 30cm tanks and fed daily with pellet food or fish that died of natural causes in the laboratory.

### Body length and weight measurements

After two weeks of acclimatization to laboratory conditions, the body length (distance from the tip of mouth to the base of caudal fin) of 32 zebrafish, 12 flying barbs and 11 whitespots was measured from photographs taken using ImageJ (Schneider et al. 2012). The fish (20 zebrafish, 22 flying barbs and 19 whitespots) were weighed to the nearest 0.1mg using a digital balance and experiments commenced 48 hours after the measurements were taken.

### Measurement of time spent foraging, proportion food intake and oxygen consumption across species

After a 24-hour period of starvation, a single species shoal comprising 5 individual per shoals (zebrafish, flying barbs or whitespots), or a mixed-species shoal (comprising 2 zebrafish, 2 flying barbs and 1 whitespot) was introduced into a 30 × 20 × 20cm glass tank. The proportion of each species in a shoal was decided based on their relative abundance in their natural habitats (pers. obs.). To simulate natural conditions where scanty or abundant food are likely to be found, freeze dried blood worms were provided in low (3 worms (≈0.7mg)) or high (30 worms (≈17.0mg)) amounts. A study by Spence et al. (2006) has shown that these species mostly reside and feed at the top of the water column.Recent stable isotope analysis studies have shown that dietary habits among surface and column feeding co-occurring species from similar aquatic habitats in West Bengal are broadly comparable (Mondal and Bhat 2021). Thus, in our experiment, we provided a single kind of food. After a 2 – minute acclimation period following the introduction of the shoal into the test tank, the food (either low or abundant food) was introduced onto the water surface, at a random corner of the tank. The shoal was recorded using a digital video camera (Canon Legria HF R306) placed directly above the test arena, for 10 minutes. From the videos, the time taken to consume 3 worms (i.e., low food treatments) or 30 worms (i.e., high food treatments) was determined and taken as the foraging time. Secondly, to check for inter-species differences in food consumption in mixed shoals, we calculated the number of bites per individual, per species. Thirdly, to check whether the foraging behavior of individual zebrafish change based on the shoal composition (i.e., single species or mixed-species), we calculated the average number of worms consumed by zebrafish individuals in single species shoals and in mixed-species shoals. In the low food condition, we tested 15 zebrafish shoals, 14 flying barb shoals, 13 whitespot shoals and 15 mixed shoals. In the high food condition, 15 zebrafish shoals, 16 flying barb shoals, 15 whitespot shoals and 15 mixed shoals were tested. Therefore, a total of 118 shoals (57 shoals in low food condition and 61 in high food condition) were tested. Fish within shoals were tested only once per food treatment condition.

Oxygen consumption of a fish is proportional to metabolism (Martins et al. 2011). In order to understand the relationship between foraging time and metabolism for each species, oxygen consumed in one hour by individuals (of different species) was measured. We measured the oxygen intake (in mg/l) from ambient water by test individuals using a DO meter and. An in-house respirometer (See A1. Measurement of oxygen consumption, and Figure S1 in Supplementary Information for details of the method).

### Shoaling preference in zebrafish in the presence of a predator

The experimental tank (30cm × 30cm × 62cm) was divided into three compartments using two transparent perforated partitions (to enable visual and olfactory cue exchange). The compartments at the two ends (11cm × 30cm × 30cm) was used for the stimulus shoals. While one stimulus shoal had 10 conspecifics (i.e., a zebrafish shoal), the other had 5 zebrafish, 4 flying barbs and 1 whitespot (i.e., a mixed shoal). The stimuli shoals were swapped every 10 trials to prevent side bias. The central compartment (40cm × 30cm × 30cm) comprised three zones: two association zones that were 2.5cm (i.e., approximately one body-length distance) away from the end chambers, were demarcated by lines drawn on the wall of the tank and a middle zone. Test individuals were considered associating with a stimulus shoal if any part of their body was inside the association zone. A transparent and perforated chamber (20cm × 30cm × 9.5cm) was attached to the wall of the middle zone. In half of the trials, this chamber contained a snakehead (*Channa punctatus*). The experimental tank was filled up with aged water to 10cm depth and the stimuli shoals and the predator (in half of the trials) were introduced. After a period of 10 minutes, a test fish was released in the middle compartment (See Figure S2, Supplementary Information for a schematic of the experimental arena). The movement of the test fish was then recorded for 10 minutes using a digital video-camera (Canon Legria HF R306) placed near the side of the experimental tank. Association preferences (i.e., time spent in the left and right association zone) of the test individual for either of the stimuli shoals were quantified by an observer blind to the treatment type. A total of 30 zebrafish subjects were tests in the presence and 30 subjects were tested in the absence of the predator.

To check whether the predator had a prey species preference, an additional set of two-choice trials were conducted where 11 *Channa punctatus* individuals (previously starved for 48 hours) were given a choice between two different prey species (zebrafish/flying barb/whitespot) placed in two opposing chambers, with the *Channa* (predator) placed in the central arena (see Supplementary Information A5 for details of the experiment).

### Shoaling preferences when the abundance of conspecifics varied in the stimuli shoals

Low abundances of conspecifics in the environment may drive species to shoal with heterospecifics (Paijmans et al., 2019). Here, by performing two-choice experiments, we investigated whether zebrafish associate with other species because of low abundance of conspecifics. The experimental arena described above (but without the predator chamber) was used to conduct the experiments (Figure S3, Supplementary Information). It is established that zebrafish prefer larger conspecific shoals over smaller conspecific shoals only when the smaller shoal have 4 or lesser number of individuals (Seguin and Gerlai 2017). Based on this, we presented test fish a choice between shoals comprising 5 individuals: a conspecific shoal comprising 5 zebrafish, and a mixed shoal comprising either 3 zebrafish, 1 flying barb, 1 whitespot or 2 zebrafish, 2 flying barbs, and 1 whitespot. Therefore, we presented test zebrafish a choice between two similarly sized shoals where the number of conspecifics differed in the range it could detect. Thirty of each kind of (i.e., a total of 60) trials were performed. Additionally, to test whether zebrafish have a preference for conspecific shoals over heterospecific shoals, we performed 10 trials in which test zebrafish were given a choice between a conspecific shoal and a heterospecific shoal (flying barb shoals) were tested.

### Familiarity as a factor in shoal association choice

Familiarity is a strong driver of shoaling decisions (Catellan et al. 2019; Barber and Wright 2001) and species have been reported to shoal with familiar heterospecifics over unfamiliar conspecifics (Ward et al. 2003). To examine the role of familiarity in shoal association preferences, we tested the choice of zebrafish individuals when presented with shoals differing in the extent of their familiarity to the test individual. In the present study, individuals were considered familiar when they belonged to the same population and had been maintained in the same stock tank for a period of at least 30 days in the laboratory. Fish raised in isolation for numerous generations are considered unfamiliar to one another (Martin et al. 2022). Based on this, ‘unfamiliar’ individuals for this experiment were collected from a similar shallow slow-moving stream (situated near Kharagpur, 180 kms away from the location of the first population at Haringhata, in West Bengal, India) and were kept in a separate stock tank. This, thus, ensured that the test zebrafish had never encountered the unfamiliar individuals. Individuals from the two populations were not significantly different in terms of size (Table S1, Supplementary Information) or any other external morphological features such as colouration or pigmentation (pers obs.).

The experimental arena for the experiment was the same as was used in the previous shoal choice experiment (Figure S3, Supplementary Information). We assembled four kinds of stimuli shoals (familiar conspecific shoals or FC shoals, familiar mixed shoals or FM shoals, unfamiliar conspecific shoals or UC shoals, unfamiliar mixed shoals or UM shoals). For each type of experimental trial, any of these two stimulus shoals were placed in the end compartments. Thus, 6 combinations of trials and, a total of 141 trials were performed. Test zebrafish individuals had to choose between: (1) familiar conspecific shoals (FC shoals) versus familiar mixed shoals (FM shoals) (n=30 trials), (2) familiar conspecific shoals (FC shoals) versus unfamiliar conspecific shoals (UC shoals) (n=10 trials), (3) familiar conspecific shoals (FC shoals) versus unfamiliar mixed shoals (UM shoals) (n=30 trials), (4) familiar mixed shoals (FM shoals) versus unfamiliar conspecific shoals (UC shoals) (n=30 trials), (5) familiar mixed shoals (FM shoals) versus unfamiliar mixed shoals, (UM shoals)(n=11 trials) and, (6) unfamiliar conspecific shoals (UC shoals) versus unfamiliar mixed shoals (UM shoals) (n=30 trials). For experiments involving testing of choice between familiar conspecific shoals (FC shoals) and unfamiliar conspecific shoals (UC shoals) fewer trials (n=10) were performed because it is well known that zebrafish prefer familiar over unfamiliar conspecifics (Gerlach and Lysiak 2006) and here intended to verify what is already well established. Further, for choice tests between familiar mixed shoals (FM shoals) and unfamiliar mixed shoals (UM shoals) some unfamiliar individuals were lost (i.e., due to death) from the other species (members of UM shoals). Thus, 11 choice test trials of this kind could be performed.

## Statistical Analysis

Statistical analyses were performed using R, version 4.0.2 (R Core Team 2020). Generalized linear models (GLM) were built (using “glm” function in R) to check: (i) the effect of species on individual specific traits (i.e., body-length, weight, oxygen consumption) and (ii) the effect of shoal composition on foraging time. Before developing the models, the distribution of the data was checked using the ‘fitdistr’ function (Delignette and Dutang, 2015). As our data was not close to any specifically defined distribution, we did not add a link function into our models. In such GLMs the individual traits (length, weight, or oxygen consumption), foraging time or time spent associating with the stimuli shoals were the dependent variable/s, and the species or treatment (shoal composition) were the fixed factor. All model comparisons were performed using ANOVA in ‘car’ package (Fox and Weisberg, 2019) and post hoc paired tests (Tukey’s post hoc HSD Test) were carried out for comparing the effects of factors that were significant. Chi square tests were performed to check whether the average food consumption by zebrafish individuals in conspecific shoals is comparable to the average food consumption by zebrafish individuals in mixed-species shoals. To compare the differences in time spent associating with mixed-species shoals in the presence and absence of a predator we performed Wilcoxon unpaired tests. Further, Wilcoxon paired tests were performed to compare association time of test zebrafish individuals to shoals differing: (i) in the abundance of conspecifics or (ii) in familiarity and species composition. The cut-off level for significance was set as p<0.05 for all comparisons.

## Results

### Body length and weight measurements

Zebrafish differed significantly from the other two species in terms of body length as well as their weights. The body length of zebrafish (Mean±S.E, 2.48±0.08cm) was significantly less than flying barbs (Mean±S.E, 3.86±0.09cm) and whitespots (Mean±S.E, 4.22±0.14cm) (GLM: n_flying barb_=13, n_zebrafish_= 32, n_whitespot_=11, Wald type IIχ 2 = 193.61, df = 2, p<0.0001; Tukey’s HSD Test results: whitespot vs. zebrafish: Z value=-12.06, p<0.0001, flying barb vs. zebrafish: Z value=9.87, p <0.0001) (GLM details in Table S2, Supplementary Information). In terms of weight, zebrafish (Mean±S.E, 0.26±0.03g) weighed significantly less than flying barbs (Mean±S.E, 0.51±0.02g), which in turn weighed significantly less than whitespots (Mean±S.E, 0.81±0.05g) (GLM: n_flying barb_=22, n_zebrafish_= 20, n_whitespot_=19, Wald type IIχ 2 = 115.22, df = 2, p<0.0001; Tukey’s HSD Test results:zebrafish vs. flying barbs: Z value=5.16, p <0.0001, zebrafish vs. whitespot: Z value=-10.73, p<0.0001, whitespot vs. flying barbs: Z value=-5.87, p<0.0001) (GLM details in Table S3, Supplementary Information).

### Measurement of time spent foraging, proportion food intake and oxygen consumption across species

In low food treatments, there was a significant effect of shoal composition on feeding time (GLM: n_zebrafish_=15 shoals, n_flying barb_= 14 shoals, n_whitepot_=13 shoals, n_mixed_=15 shoals, Wald type IIχ 2 = 13.89, df = 3, p<0.01; Table 1a). The feeding time of zebrafish shoals (Mean±S.E, 5.06±1.28s) and flying barb shoals (Mean±S.E, 6.85±0.89s) was significantly lower than whitespot shoals (Mean±S.E., 13±2.13s) (Tukey’s HSD Test results: zebrafish vs. whitespot: Z value= -3.58, p<0.01; flying barb vs. whitespot: Z-value= -2.72, p=0.03). The feeding time of mixed shoals (Mean±S.E., 8.66±1.51s) was comparable to that of shoals comprising a single species (zebrafish shoals: Mean±S.E, 5.06±1.28s; flying barb shoals: Mean±S.E, 6.85±0.89s; whitespot shoals: Mean±S.E, 13±2.13s) (Full details of Tukey’s HSD Test results in Table S4, Supplementary Information) (Figure 1a).

**Table 1 a).** Results of the generalized linear model (GLM) for predicting effect of species on foraging time under low food conditions

|  | Estimate | Std. Error | t value | Pr(>\|t\|) |
| --- | --- | --- | --- | --- |
| (Intercept) | 13.00 | 1.62 | 8.01 | <0.0001 |
| Zebrafish | -7.93 | 2.21 | -3.58 | <0.001 |
| Mixed | -4.33 | 2.21 | -1.95 | 0.75 |
| Flying barb | -6.14 | 2.25 | -2.72 | <0.0001 |

Model: Foraging time ~ Species
Coefficients:

**Figure 1:**
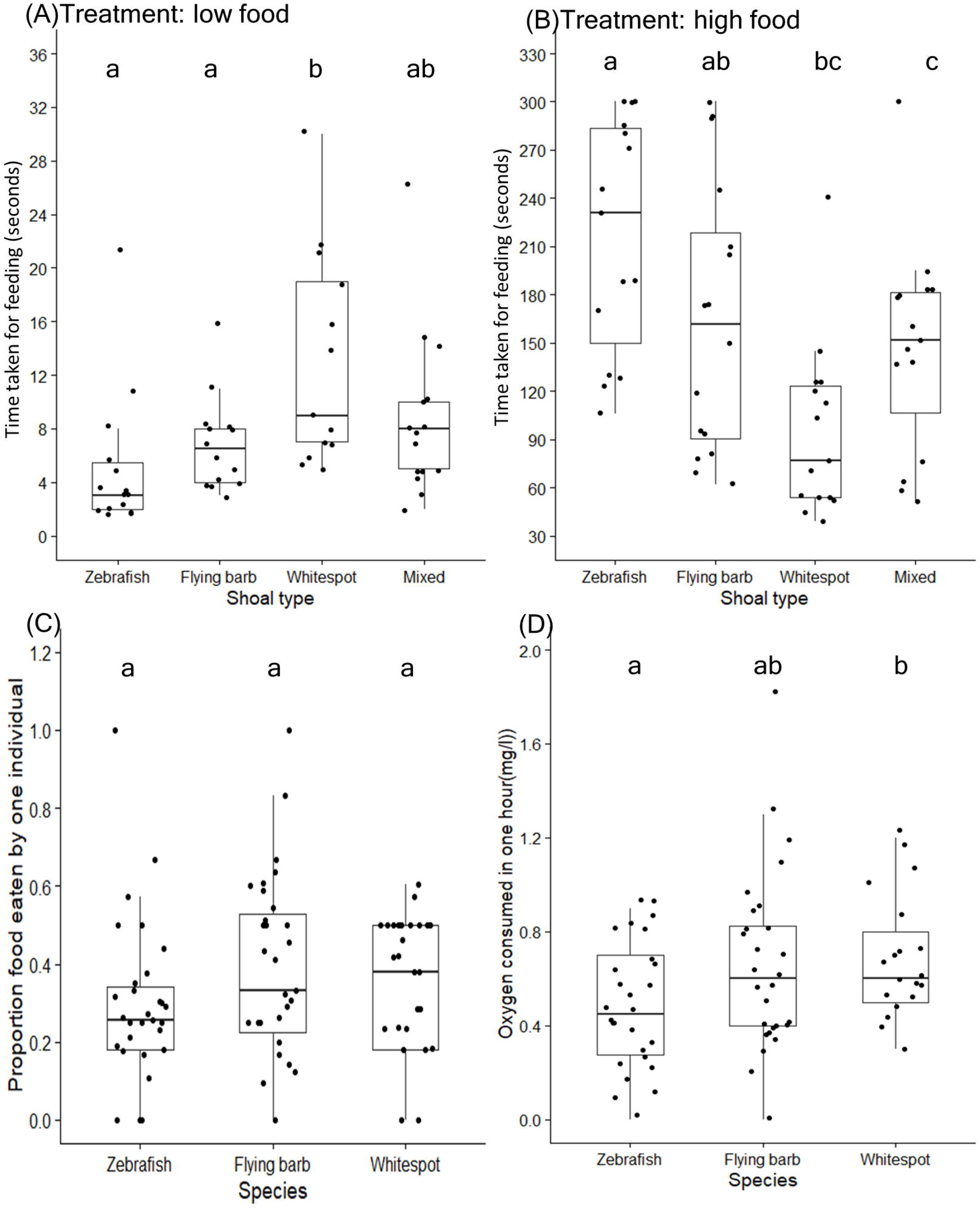
Time spent foraging, proportion food intake and oxygen consumption across species. Time taken for feeding (in seconds) in (A) low food treatments and (B) high food treatments. (C)The proportion food eaten per individual for different species within mixed shoals and (D) the oxygen consumption in one hour (mg/l) by individuals belonging to species. Data points are represented as dots. The different letters placed above the boxes represent significant differences between the categories. The comparisons have been made by performing GLM followed by Tukey’s HSD Test (p<0.05).

In high food treatments, there was a significant effect of shoal composition on feeding time (GLM: n_zebrafish_=n_whitepot_=n_mixed_=15 shoals, n_flying barb_=16 shoals, Wald type IIχ 2 = 23.11, df = 3, p<0.0001; Table 1b). In contrast to low food treatments, the feeding time of whitespot shoals (Mean±S.E., 94.8±16.3s) was significantly lower than that of zebrafish (Mean±S.E., 216.6±18.12s) and flying barb shoals (Mean±S.E., 164.75±20.4) in the presence of high food (Tukey’s HSD Test results: whitespot vs. zebrafish: Z value= 4.75, p<0.001; whitespot vs. flying barb: Z value= 2.77, p=0.02). Under high food condition, the feeding time of mixed shoals (Mean±S.E., 146.8±16.3s) was significantly lower than zebrafish shoals (Mean±S.E., 216.6±18.12s) (Tukey’s HSD Test results: mixed shoal vs. zebrafish shoal: Z value= -2.72, p=0.03) and, the feeding time of mixed shoals (Mean±S.E.,146.8±16.3s) was comparable to that of whitespot (Mean±S.E., 94.8±16.3s) and flying barb shoals (Mean±S.E., 164.75±20.4) (Full details of Tukey’s HSD Test results: Table S5, Supplementary Information) (Figure 1b).

**Table 1 b).**
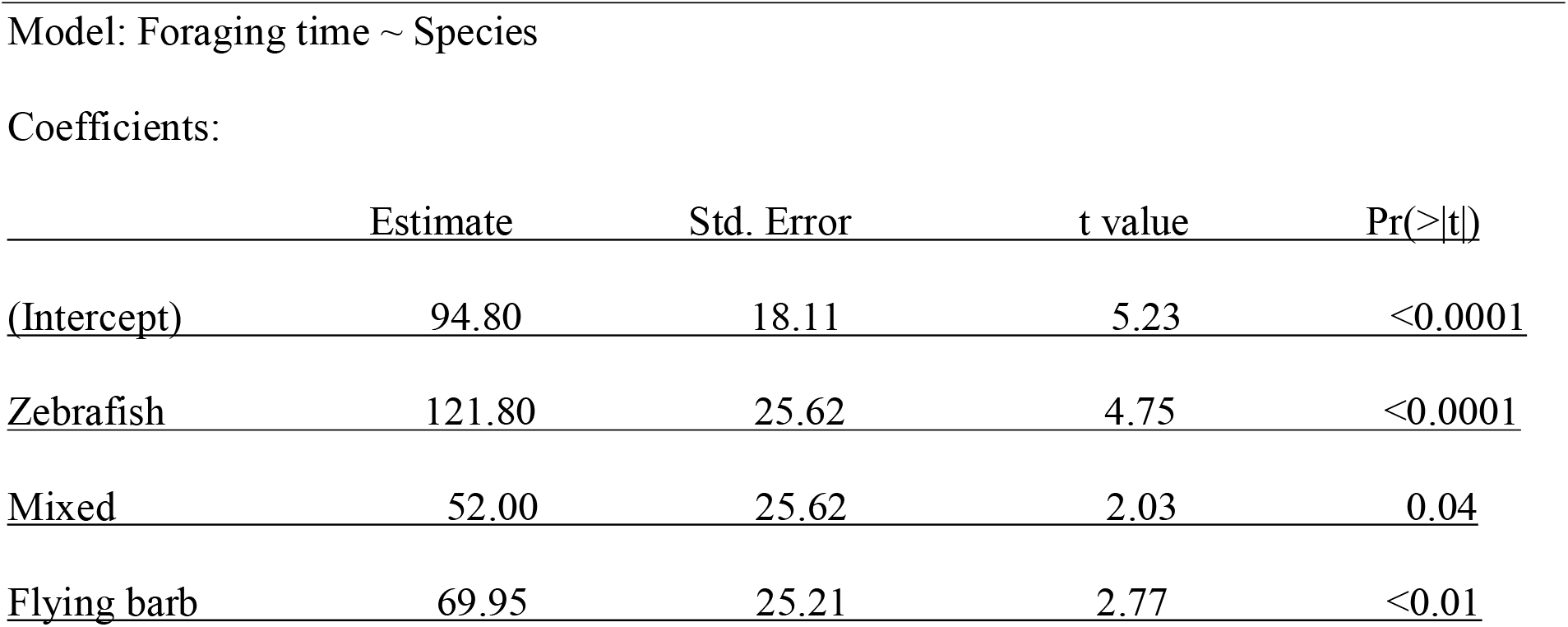
Results of the generalized linear model (GLM) for predicting effect of species on foraging time under high food condtitions

|  | Estimate | Std. Error | t value | Pr(>\|t\|) |
| --- | --- | --- | --- | --- |
| (Intercept) | 94.80 | 18.11 | 5.23 | <0.0001 |
| Zebrafish | 121.80 | 25.62 | 4.75 | <0.0001 |
| Mixed | 52.00 | 25.62 | 2.03 | 0.04 |
| Flying barb | 69.95 | 25.21 | 2.77 | <0.01 |

Within mixed shoals, the proportion of food eaten per individual per species was comparable across all three species (GLM: n=15; Wald type IIχ 2 = 3.11, df = 2, p=0.21) and treatment types i.e., high food or low food conditions (GLM: Wald type IIχ 2 = 0.02, df = 1, p=0.87; Table 1c). (Figure 1c; Table 1c). Further, the average food consumption by zebrafish individuals was comparable when they were in conspecific shoals and in mixed-species shoals (Chi square test results for low food treatments: χ^2^= 0.01, df = 2, p = 0.99; Chi square test results for high food treatments χ^2^ = 0.22, df = 2, p = 0.89).

**Table 1 c).** Results of the generalized linear model (GLM) for predicting effect of species and treatment (high food or low food) on proportion food intake by individuals.

|  | Estimate | Std. Error | t value | Pr(>\|t\|) |
| --- | --- | --- | --- | --- |
| (Intercept) | 0.34 | 0.04 | 7.57 | <0.0001 |
| Zebrafish | -0.05 | 0.05 | -0.98 | 0.32 |
| Flying barb | 0.04 | 0.05 | 0.77 | 0.43 |
| Low food | -0.01 | 0.04 | -0.15 | 0.87 |

Coefficients:

The oxygen consumption levels across the three species were then compared. Among the three species, there was a significant difference in the oxygen consumption levels between zebrafish (Mean±S.E., 0.46±0.05mgl^-1^hour^-1^) and whitespot (Mean±S.E., 0.69±0.05 mgl^-1^hour^-1^) (GLM: n_zebrafish_=n_flying barb_= 28, n_whitespot_=19, Wald type IIχ 2 = 8.31, df = 2, p=0.01, Tukey’s HSD Test results: whitespot vs. zebrafish: Z value= -2.58, p=0.02). The oxygen consumption in flying barbs (Mean±S.E., 0.66±0.06 mgl-1hour-1) were intermediate to the other two species (i.e., zebrafish and white spots) and were not significantly different from either of them. There was no significant difference between the methods of measurement (GLM: n_respirometer_= 24, n_DOmeter_=51, Wald type IIχ 2 = 2.48, df = 1, p=0.11) (Figure 1d; Table 1d).

**Table 1 d).** Results of the generalized linear model (GLM) for predicting effect of species and method (DO meter or in-house respirometer) on oxygen consumption by individuals.

|  | Estimate | Std. Error | t value | Pr(>\|t\|) |
| --- | --- | --- | --- | --- |
| (Intercept) | 0.74 | 0.07 | 9.4 | <0.0001 |
| Zebrafish | -0.24 | 0.09 | -2.58 | 0.01 |
| Flying barb | -0.05 | 0.09 | -0.53 | 0.59 |
| Low food | -0.12 | 0.07 | -1.57 | 0.11 |

### Shoaling preference in presence of a predator

Test zebrafish associated with a mixed shoal significantly more in the presence of a predator (Mean ±S.E., 297.2±33.61s) than in absence of a predator (Mean ±S.E., 169±23.19s) (Wilcoxon Unpaired test results: W=270.5, n=30, p<0.01) (Figure 2). Further, in treatments where the predator was present, the number of visits by test individuals to the mixed shoal was comparable between the first two minutes (Mean ±S.E., 2.66±0.48) and last two minutes (Mean ±S.E., 2.73±0.38) of the recording (V = 108.5, n=30, p = 0.81). We also conducted experiments where test predators (snakehead individuals) were given a choice between a zebrafish shoal and a flying barb shoal. Although test snakeheads struck comparably at the zebrafish shoal (Mean±S.E., 4.72±1.06 times) and flying barb shoal (Mean±S.E., 4.90±1.06 times) (Wilcoxon paired test results: V=28, n=11, p = 1), the first two strikes were significantly more towards flying barb shoals (towards zebrafish shoal: Mean±S.E., 0.45±0.19 times; towards flying barb shoal: Mean±S.E., 1.54±0.19 times) (Wilcoxon paired test results: V=4.5, n=11, p =0.04).

**Figure 2:**
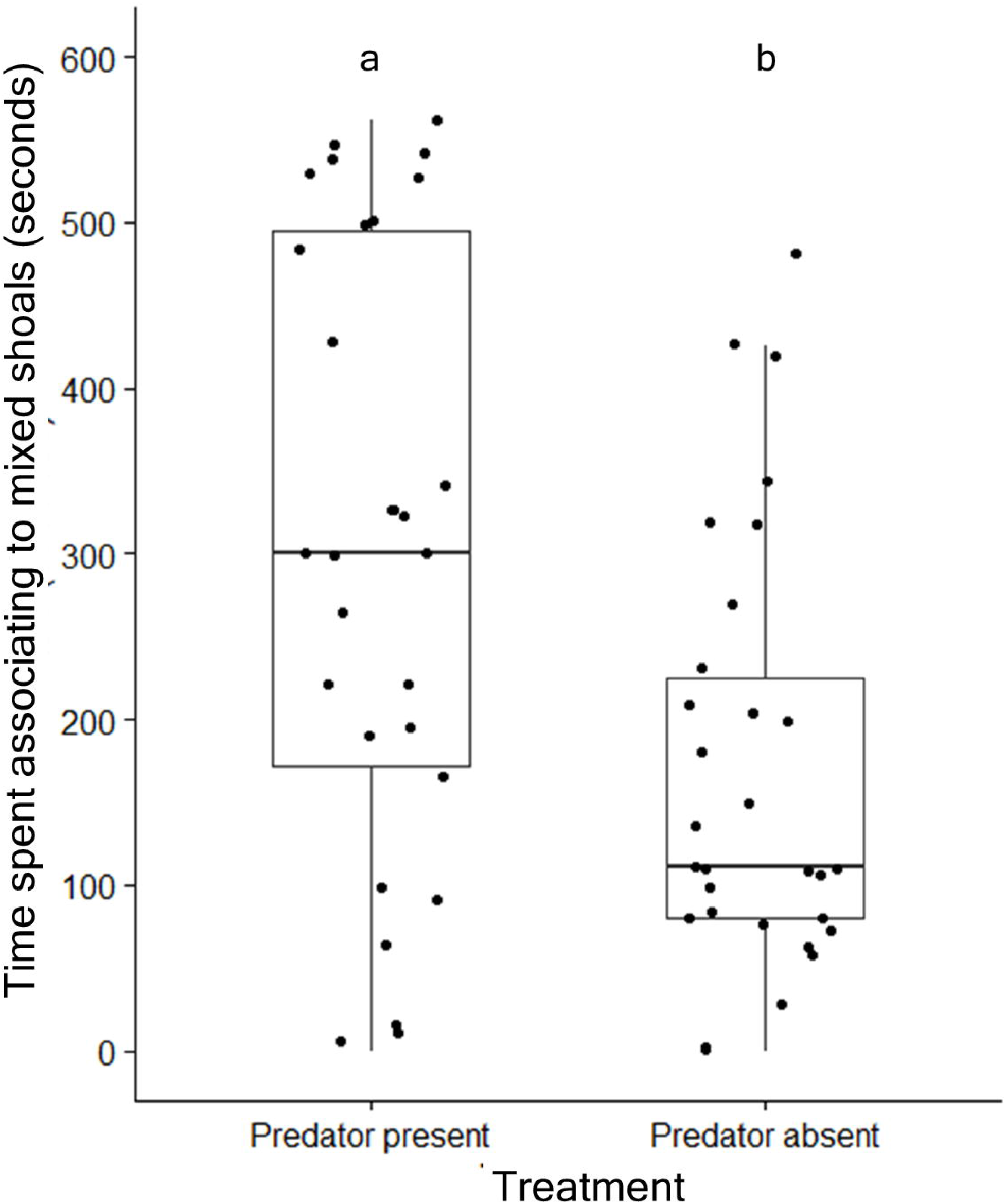
Time spent by test zebrafish in association with mixed shoals, in the presence and absence of a predator. Data points are represented as dots. The different letters placed above the boxes represent significant differences between the categories. The comparisons have been made by performing Wilcoxon unpaired test (p<0.05).

### Shoaling preference when the abundance of conspecifics varied

Test zebrafish showed similar preference for a conspecific shoal and a mixed shoal comprising different proportions of conspecifics, where the overall number of individuals in both shoals remained same. The association time with a conspecific shoal comprising five zebrafish (Mean ±S.E., 252±30.44s) was comparable to association time with a mixed shoal comprising two zebrafish, two flying barbs and one whitespot (Mean ±S.E., 174.5±26.45s) (Wilcoxon paired test results, V = 295, n=30, p = 0.21, Figure 3a). Similarly, the association time with a conspecific shoal comprising five zebrafish (Mean ±S.E., 197.13±27.87s) was comparable to association time with a mixed shoal comprising three zebrafish, one flying barb and one whitespot (Mean ±S.E., 178.13±24.22s) (Wilcoxon paired test results: W = 249.5, n=30, p = 0.74), Figure 3b). We found that test zebrafish individuals showed a preference for a conspecific shoal over heterospecific shoal (i.e., with only flying barbs and no zebrafish) (Wilcoxon paired test results: V=50.5, n=10, p=0.02).

**Figure 3:**
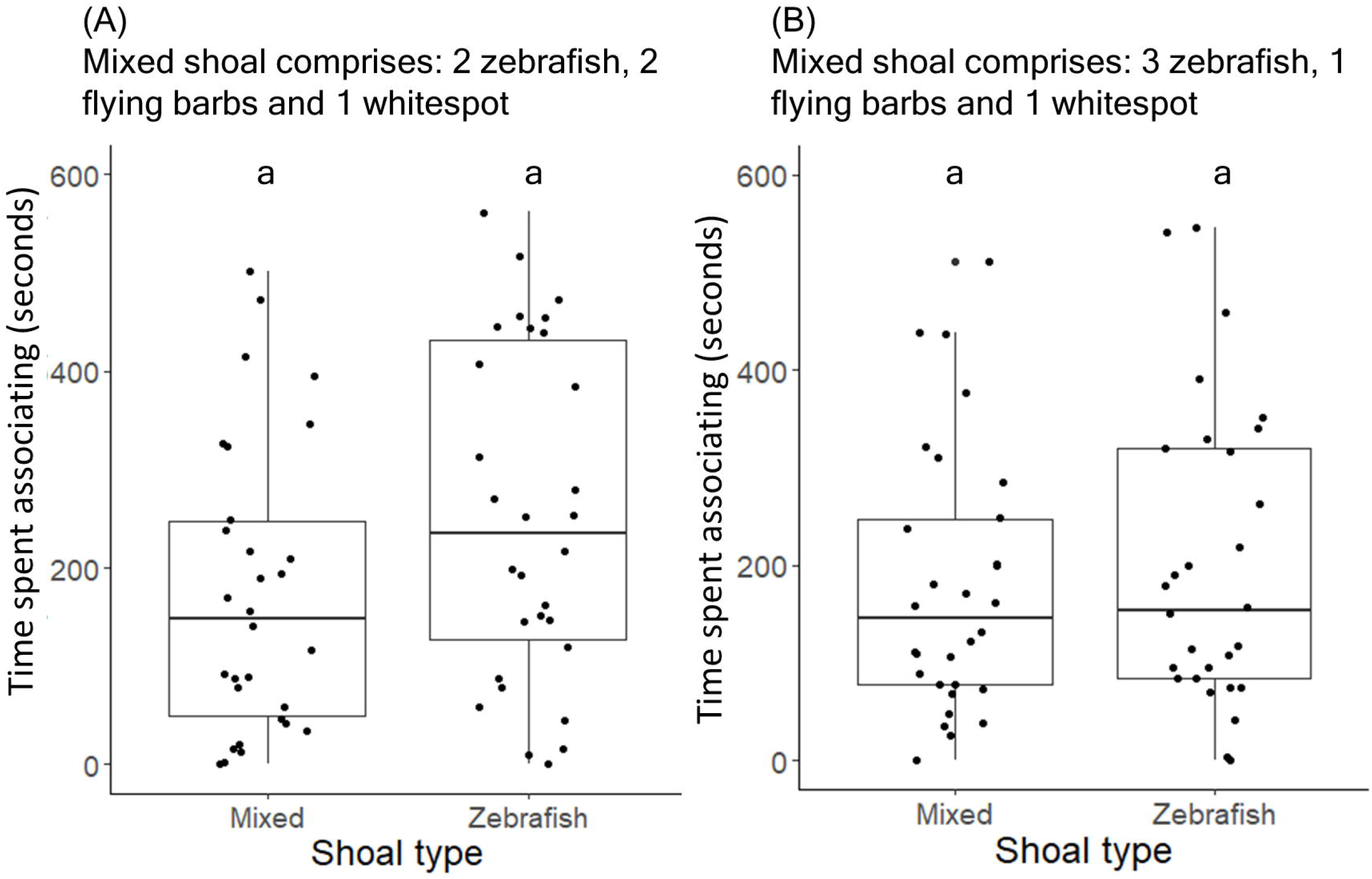
**Time spent by test zebrafish in association with: (A) conspecific shoals comprising 5 zebrafish and mixed shoals comprising 2 zebrafish, 2 flying barbs and 1 whitespots and (B) conspecific shoals comprising 5 zebrafish and mixed shoals comprising 3 zebrafish, 1 flying barb and 1 whitespot**. Data points are represented as dots. The different letters placed above the boxes represent significant differences between the categories. The comparisons have been made by performing Wilcoxon unpaired test (p<0.05).

### Familiarity as a factor in shoal association choice

When presented a choice between shoals varying in familiarity and species composition (i.e., conspecific shoals and mixed shoals), test zebrafish had a greater association time with FC shoals (Mean ±S.E., 273.93±27.91) than UM (Mean ±S.E., 150.66±21.33) (Wilcoxon paired test results: FC vs. UM: V=343, n=30, p=0.02) and FC shoals (Mean ±S.E., 241.9±40.96) than UC shoals (Mean ±S.E., 64.5±10.97) (Wilcoxon paired test results: V = 3, n=10, p= 0.02). Test zebrafish showed comparable associations with stimuli shoals in the other choices provided (Wilcoxon paired test results: FM vs. UC: V = 238, n=30, p = 0.91; UM vs. UC: V=212, n=30, p= 0.68; FC vs. FM: V=256, n=30, p=0.64; FM vs. UM: V = 27, n=11, p = 0.62).

## Discussion

The main aim of the study was to unravel the multiple roles of the presence of conspecific and co-occurring species, familiarity, and predation risk on the shoaling decisions among zebrafish. For this, we used a multipronged approach based on a variety of choice-based experiments to specifically understand the effect of these factors on shoal choice. Firstly, to investigate whether forming mixed-species shoals benefit zebrafish, we tested individuals in four different contexts. Our results indicate that food availability, protection from predators and kin familiarity together drive zebrafish towards forming shoals with other species. While overall feeding time varied depending on the species composition of the mixed-species shoals, each species consumed comparable amount of food. Shoal choice experiments revealed that zebrafish prefer a mixed shoal under predation threat and familiar shoals over unfamiliar shoals. Interestingly, the abundance of conspecifics did not impact shoaling decisions. These results indicate that there are certain advantages to forming mixed-species shoals. We speculate that, the above benefits, combined with a variety of factors may provide an explanation for the formation of mixed-species shoaling decisions in zebrafish.

### Measurement of time spent foraging, proportion food intake and oxygen consumption across species

Zebrafish are significantly smaller (i.e., have lower total body length and lower weight) than the other two species and have a lower metabolic rate than whitespots (as it consumes significantly less oxygen than the whitespots). The morphological features of the mouth and foraging microhabitat of these species have been characterized in earlier studies: flying barbs and zebrafish have mouths obliquely directed upwards with a longer lower jaw and, are surface as well as column feeders (Das et al. 2010; Spence et al. 2006). Whitespots, on the other hand, have a small protrusible mouth (pers. obs.) and are chiefly surface feeders (Spence et al. 2006). In addition to mouth parts and feeding habits, our studies indicate that the oxygen intake is also comparable between flying barbs and zebrafish, and that of whitespots is significantly greater. We speculate that similarities (or differences) in time spent foraging is a result of similarities in size, mouth types, feeding habits and oxygen intake (thus, metabolic rate). The basic premise of the optimal foraging theory states that food intake increases linearly with size (Werner 1974). However, in our study, despite differences in size (body length and weight) between zebrafish and flying barbs, the species were found to take similar amounts of time to consume equal amount of food. It is possible that had body-size differences between the species been greater, it could translate to differences in foraging time.

Interestingly, we found that the pattern for time spent foraging reversed when food quantity was switched from low to high. Under low food conditions, as reflected by the low foraging time, zebrafish and flying barbs consumed food faster than whitespots. It is likely that under high food conditions, zebrafish and flying barbs got satiated and thus shoals had a longer feeding time. We further suggest that in this experimental treatment, whitespots (being larger and having a greater metabolic rate) were not satiated and had a significantly lower feeding time than the other two species. The foraging time for mixed shoals was comparable and intermediate to that of single species shoals. This suggests that even in mixed-species shoals, individuals did not deviate from their foraging patterns. Similar to our findings, other experiments involving *Pseudomugil signifier* and *Gambusia holbrooki* show that foraging behavior is not plastic, and species are unable to adjust foraging behavior in mixed-species shoals (Keiller et al. 2021).

Despite differences in foraging time, inter-species food apportionment was observed in mixed-species shoals. There are two likely reasons for these. Firstly, differences in aggression and foraging behavior across the species might be a driver of mixed-species shoaling. Whitespots, the largest of the species are believed to show non-aggressive behavior towards other species and co-exist as mixed shoals (Daniels 2002). Secondly, tropical ponds and ditches are abundant in phytoplankton and thus, food sources being abundant, competition is unlikely to be high in mixed shoals. In contrast to our findings, inter-species competition for food resulting in unequal foraging amongst species often occur in mixed shoals (Mathis and Chivers 2003; Lukoschek and Mccormick 2000; Paijmans et al.,2020; Camacho-Cervantes et al. 2019). In our study, comparable amount of food was consumed by zebrafish individuals in single species shoals and as mixed-species shoals. This suggests that interspecies competition for food is absent within these tropical shoals comprising zebrafish, flying barbs and whitespots. Thus, low or near absent inter-species competition is an important likely factor that keeps the species together. Interestingly, despite being significantly smaller than both species and having a lower metabolic rate than whitespots, zebrafish consume comparable amount of food as the other two species. Therefore, foraging benefits in the form of equal access to food could be another driver for zebrafish to form mixed-species shoals.

### Shoaling preference in presence of a predator

Apart from differences in body size, the three fish species also differ in their external morphology/appearance. Zebrafish have five uniformly pigmented horizontal stripes, while flying barbs have a single dark, broad lateral line (sometimes absent) and whitespots lack stripes but have a dark blotch at the lower third of the dorsal fin (Das et al. 2010). Interestingly, results on shoal choice experiments in the presence of a predator indicated that, test zebrafish spent significantly more time with a mixed shoal (comprising dissimilar individuals) as compared to a single species (heterospecific) shoal. Thus, dissimilarity in size and phenotype does not play a role in shoaling decisions under predator threat i.e., oddity does not control shoaling decision. In contrast to our findings, size segregation in shoals (to minimize oddity) often occur in the presence of a predator: mixed cyprinid shoals and shoals comprising certain minnow species segregate in accordance with size in the presence of a predator (Hoare et al. 2000; Allan and Pitcher 1986; Theodrakis1985). In laboratory tests where the predator was given a choice between a flying barb shoal and a zebrafish shoal, starved *Channa* individuals were found to strike at flying barbs shoal more often than zebrafish shoals. Therefore, we speculate that the preferred association of test zebrafish to mixed shoals could be due to the predators’ preference for whitespots and flying barbs over zebrafish. Across taxa, predators selectively favour prey species based on size (body length) and body mass (Owen-Smith and Mills 2008; Dallal et al. 2019; Prado et al. 2020). Flying barbs and whitespots are larger in size and slower swimmers as compared to zebrafish (pers. obs.), and thus, for an ambush predator like the *Channa*, catching these species is more beneficial (in terms of net energy gained). Armored brook spined sticklebacks (*Culaea inconstans*) associated more with non-armored minnow shoals (*Pimephales promelas*) than with conspecific shoals. This is because the predator (perch) selectively attacked minnows in mixed shoals, thereby reducing the vulnerability of the three spined sticklebacks (Mathis and Chivers 2003). Apart from differential preference by the predator, mixed shoals might also provide higher vigilance (using heterospecific alarm cues) and could be an additional reason for mixed shoaling in the presence of predators (Ward et al. 2018). In consensus with our findings, damselfish (*Abudefduf vaigiensis*) associate equally with conspecific shoals and shoals of the phenotypically different Australian mado (*Atypichthys strigatus*), indicating that oddity is not a universal driver of shoaling decisions (Paijmans 2020).

### Shoaling preference when the proportion of conspecifics varied

Contrary to expectations, our results indicated that the abundance of species have no impact on shoaling decisions: individuals show equal preference to shoals differing in proportion of conspecifics. In agreement with other studies (Snekser et al. 2010; Saverino and Gerlai 2008), we found that test zebrafish individuals showed a preference for conspecific shoals over heterospecific shoals (where only flying barbs were present). Shoaling with other species have disadvantages such as reduced mating opportunities and increased oddity. Yet, test zebrafish associate comparably with mixed-species shoals and conspecific shoals. We speculate that advantages of shoaling with other species is comparable to the disadvantages (reduced mating opportunities, increased oddity), and thus, zebrafish make shoaling decisions irrespective of abundance of conspecifics. Another possibility is that in natural environments, zebrafish are unable to distinguish heterospecifics from conspecifics and thus, shoals form when two or more individuals are in spatial proximity to one another. In natural environments, under turbid conditions and low visibility, visual cues may not be efficient. Chubs (*Leuciscus cephalus*) rely greatly on olfactory cues to distinguish conspecifics from heterospecifics (Ward et al. 2019). We speculate that in environments with water flow and turbidity, detection and differentiation of olfactory cues between co-occurring species might be harder. This might be an explanation for similar preference of test zebrafish for conspecific shoals and mixed-species shoals. Further studies testing sensory mechanisms in recognizing heterospecifics and conspecifics will provide some insight into the mechanisms of forming mixed-species shoals.

### Familiarity as a factor in shoal association choice

The test zebrafish showed comparable association between conspecific and mixed shoals (when stimulus shoals are either familiar or unfamiliar i.e., FM vs. FC and UM vs. UC). We speculate that this occurs because associating with either shoal can be beneficial: associating with mixed shoals have benefits such as improved predator avoidance and food apportionment (as discussed in the previous sections). Associating with conspecific shoals can provide mating benefits, and better social communication. Further, individuals also showed comparable association between UC and FM shoals, and UM and FM shoals. We speculate that this occurs because shoaling with either shoal has an advantage. In contrast to our findings on comparable association between UC and FM shoals, test chub (*Leuciscus cephalus*) individuals preferred familiar heterospecific (minnow or *Phoxinus phoxinus*) shoals over unfamiliar conspecific shoals (Ward et al. 2003). Shoaling with mixed shoals confer anti-predator and foraging benefits. Shoaling with familiar individuals is advantageous because of better predator evasion and foraging (Griffiths et al. 2004), more stable dominance hierarchies (Höjesjö et al. 1998; Gómez-Laplaza 2005) and quicker recovery from fear (Mathuru et al. 2017). Thus, across species, fishes show a preference for shoaling with familiar individuals (Binoy and Thomas 2006; Brown 2002; Ward et al. 2003) over unfamiliar shoals (Brown and Smith 1994; Griffiths and Magurran 1997). In our choice tests too, to infer FC vs. UC preference among zebrafish individuals, we found that zebrafish associated significantly more with familiar conspecific shoals over unfamiliar ones. Test individuals also associate with FC shoals significantly more than UM shoals. We speculate that this is because combined disadvantages associated with unfamiliarity and mixed shoaling (decreased mating opportunities and unstable social hierarchies) is sufficient to drive association tendencies towards FC shoals. Across taxa, literature on mixed-species groups is scanty (Paijmans 2019) and while most studies focus on whether benefits related to either foraging or predation drive mixed-species shoaling, ours show how various ecological factors control shoaling decisions differently, thereby giving greater insight into the ultimate causes of shoaling among wild zebrafish. In addition to foraging and anti-predatory benefits and familiarity mixed shoaling can also be determined by other factors such as the inability to distinguish conspecifics and heterospecific, differences in social learning amongst the species (Paijmans 2019) and passive mechanisms such as comparable swimming speeds (Krause et al., 2005). While our study focuses on benefits obtained by zebrafish, it would be interesting to conduct investigations on what drives flying barbs and whitespots towards mixed-species shoaling. We suggest that such advantages combined with a variety of ecological factors provide ultimate explanations for the existence of such mixed-species shoals in these habitats.

## Acknowledgements

The authors thank the Indian Institute of Science Education and Research Kolkata (IISER Kolkata), India, for providing infrastructural and financial support. The authors thank Mr. Asutosh Mondal for his help in the collection of wild shoals from Haringhata. The authors also thank Prasenjit Pan for help in fish maintenance in the laboratory. IM was supported by a junior research fellowship from IISER Kolkata. This work was supported by the Academic Research Funds provided by Indian Institute of Science Education and Research Kolkata (IISER Kolkata), India to AB.

## Ethics Statement

Guidelines outlined by the Committee for the Purpose of Control and Supervision of Experiments on Animals (CPCSEA), Ministry of Fisheries, Animal Husbandry and Dairying, Government of India were followed in all aspects of maintenance and experimentation. All experimental protocols followed here have been approved by the Institutional Animal Ethics Committee’s (IAEC) and guidelines of Indian Institute of Science Education and Research (IISER) Kolkata, Government of India (Approval number IISERK/IAEC/AP/2021/70).

## Author Contributions

A.B. and I.M. conceived and designed the study. I.M. carried out experimental work behavioral assays, statistical analysis, and wrote the first draft of the manuscript. A.B. helped to significantly revise the manuscript. Both authors gave final approval for publication.

## Competing Interests

The authors declare no competing or financial interests.

## Notes

### Competing Interest Statement

The authors have declared no competing interest.

