## Supplementary Files for "What drives mixed-species shoaling among wild zebrafish? Role of predators, food access, abundance of conspecifics and kin familiarity"

Following Bolbuc et al. (2002), a flow through respirometer set up was built to measure the oxygen consumption of individuals. The chamber containing the test fish was fashioned from a clear plastic container (Volume=200ml). An oxygen probe (Hanna HI 9142 Dissolved Oxygen Meter) was connected to this chamber and this measured the dissolved oxygen concentration in the chamber. Water was constantly circulated by an electrical pump. To ensure no leakage of water from the system, water proof sealing was applied between the container and piping (Figure S1). Test fish were starved for 24 hours and were in a postabsorptive digestive state. On the day of oxygen measurement, a test fish was introduced in the clear chamber and air bubbles were gently removed from the system. Oxygen consumption readings were taken every 15 minutes for an hour.

Additionally, oxygen consumption was also measured by a introducing an oxygen probe in an airtight container (Volume=250ml) in which a test fish was present. In this set up not pump was connected and thus, water flow was absent. These measurements were also conducted every 15 minutes for an hour. While the above respirometer is a more sophisticated set up, this simple method to estimates differences in oxygen consumption across individuals.

**Figure S1**

CLEAR PLASTIC CONTAINER CONTAINING TEST FISH

RUBBER TUBING

AIR TIGHT SEALING

OXYGEN PROBE

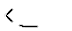

WATER CIRCULATION

PUMP

PLASTIC TUB

PROBE READING

**Figure S1:** Overhead view of the respirometer set up (not to scale).

**Figure S2**

.

5cm

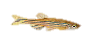

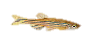

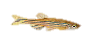

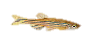

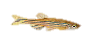

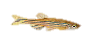

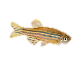

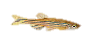

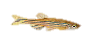

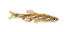

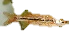

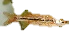

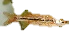

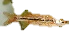

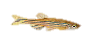

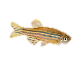

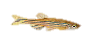

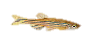

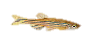

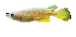

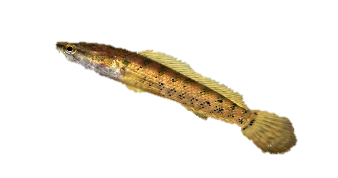

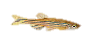

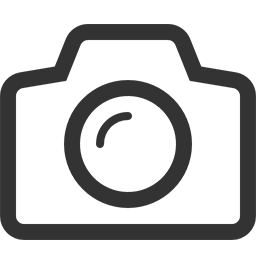

11cm

11cm

30cm

20cm

30cm

Camera

9.5cm

2.5cm

A: END COMPARTMENT HOLDING STIMULUS SHOALS

B: ASSOCIATION ZONE

C: CENTRAL COMPARTMENT

D: PREDATOR CHAMBER

A

C

B

D

A

B

**Figure S2:** Side view of the experiment tank (to scale).

**Figure S3**

**Figure S3:** Side view of the experiment tank (to scale).

A: END COMPARTMENT HOLDING STIMULUS SHOALS

B: ASSOCIATION ZONE

C: CENTRAL COMPARTMENT

.

5cm

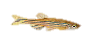

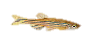

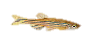

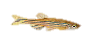

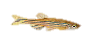

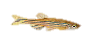

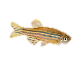

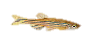

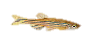

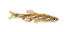

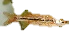

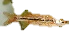

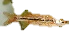

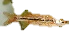

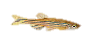

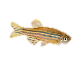

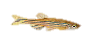

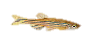

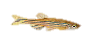

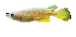

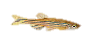

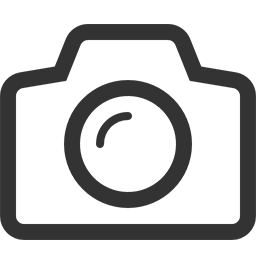

11cm

11cm

30cm

30cm

Camera

2.5cm

A

C

B

A

B

**A2. Shoaling preferences when the abundance of conspecifics varied in the stimuli shoals**

Shoals comprising 10 zebrafish conspecifics, mixed shoals comprising 7 zebrafish, 2 flying barbs, 1 whitespot and mixed shoals comprising 5 zebrafish, 2 flying barbs and 1 whitespot were assembled. In the experimental tank, one of the end-compartment housed conspecific stimuli shoal while the other end-compartment housed one of the two mixed species shoals. 10 minutes after placing the stimuli shoals, a test fish was released into the middle compartment. The shoal association preference of the test fish was observed in this period using the same protocol as mentioned in the experiment with the smaller stimuli shoals.

**A3. Comparison of size between fish from Haringhata and fish from Kharagpur (used as stimuli shoals)**

The following table summarizes body length of fish between the two populations:

**Table S1**

|  | Mean ± S.D. of body length of fish from Haringhata | Mean ± S.D. of body length of fish from Kharagpur | Wilcoxon unpaired test results |
| --- | --- | --- | --- |
| Zebrafish | 2.48±0.41 cm | 2.23±0.31 | W = 91, p-= 0.21 |
| Flying barbs | 3.86±0.28cm | 3.96±0.42 | W = 59.5, p = 0.39 |

As only one whitespot individual (from Kharagpur) was used in the unfamiliar stimulus shoal, no test was conducted to compare whitespot sizes between the two populations.

**A4. Additional results:**

*Body length and weight measurements*

**Table S2**

Results of the generalized linear model (GLM) for predicting the effect of weight on association time of test fish

Model: Weight ~Species

Coefficients:

Estimate	Std. Error	t value	Pr(>|t|)

(Intercept)	0.81	0.04	22.10	<0.0001

Zebrafish	-0.55	0.05	-10.73	<0.001

Flying barb	-0.29	0.05	-5.87	<0.001

*Measurement of feeding time*

**Table S3**

Results of the generalized linear model (GLM) for predicting the effect of shoal composition on feeding time for low food treatments

Model: Feeding time ~Shoal composition

Coefficients:

Estimate	Std. Error	t value	Pr(>|t|)

(Intercept)	13.00	1.62	8.01	<0.0001

Zebrafish	-7.93	2.21	-3.58	<0.0001

Mixed	-4.33	2.21	-1.95	0.05

Flying barb	-6.14	2.25	-2.72	0.01

*Measurement of feeding time*

**Table S4**

Tukey’s HSD Test Results for estimating differences in feeding time between shoal types (i.e. zebrafish shoals, flying barb shoals, whitespot shoals and mixed shoals) under low food conditions

Comparison	Z value	Pr(>|t|)

Mixed shoals Vs whitespot shoals	-1.95	0.20

Mixed shoals Vs zebrafish shoals	1.68	0.33

Mixed shoals Vs flying barb shoals	-0.83	0.83

Zebrafish shoals Vs flying barb shoals	0.82	0.84

Zebrafish shoals Vs whitespot shoals	-3.58	<0.01

Flying barb shoals Vs whitespot shoals	-2.72	0.03

**Table S5**

Tukey’s HSD Test Results for estimating differences in feeding time between shoal types (i.e. zebrafish shoals, flying barb shoals, whitespot shoals and mixed shoals) under high food conditions

Comparison	Z value	Pr(>|t|)

Mixed shoals Vs whitespot shoals	2.03	0.17

Mixed shoals Vs zebrafish shoals	-2.72	0.03

Mixed shoals Vs flying barb shoals	0.71	0.89

Zebrafish shoals Vs flying barb shoals	-2.05	0.16

Zebrafish shoals Vs whitespot shoals	4.75	<0.001

Flying barb shoals Vs whitespot shoals	2.77	0.02

**A5. Experiment to check snakehead’s preference towards shoals differing in species composition**

**Methods**

A 20×20×30cm^3^ bare glass tank was filled with aged water up to 12cm. Two clear perforated containers containing either a shoal comprising 4 zebrafish or a shoal comprising 4 flying barbs were introduced in the glass tank and allowed to acclimatize for 10 minutes. A 48-hour starved test *Channa punctatus* (snakehead) individual was added in the tank. The number of strikes made by the snakehead towards either shoal within 20 minutes of introduction was noted down. A total of 11 individuals (Weight, Mean±SE= 14.13±2.63) were tested.

**Results**

Although test snakeheads struck comparably at the zebrafish shoal (Mean±S.E., 4.72±1.06 times) and flying barb shoal (Mean±S.E., 4.90±1.06 times) (Wilcoxon paired test results: V=28 , n=11, p = 1), the first two strikes were significantly more towards flying barb shoals (towards zebrafish shoal : Mean±S.E., 0.45±0.19 times; towards flying barb shoal : Mean±S.E., 1.54±0.19 times) (Wilcoxon paired test results: V=4.5, n=11, p =0.04).
